# Laboratory evolution of *E. coli* with a natural vitamin B_12_ analog reveals roles for cobamide uptake and adenosylation in methionine synthase-dependent growth

**DOI:** 10.1101/2024.01.04.574217

**Authors:** Kenny C. Mok, Zachary F. Hallberg, Rebecca R. Procknow, Michiko E. Taga

## Abstract

Bacteria encounter chemically similar nutrients in their environment that impact their growth in distinct ways. Among such nutrients are cobamides, the structurally diverse family of cofactors related to vitamin B_12_ (cobalamin), which function as cofactors for diverse metabolic processes. Given that different environments contain varying abundances of different cobamides, bacteria are likely to encounter cobamides that enable them to grow robustly as well as those that do not function efficiently for their metabolism. Here, we performed a laboratory evolution of a cobamide-dependent strain of *Escherichia coli* with pseudocobalamin (pCbl), a cobamide that *E. coli* uses less effectively than cobalamin for MetH-dependent methionine synthesis, to identify genetic adaptations that lead to improved growth with less-preferred cobamides. After propagating and sequencing nine independent lines and validating the results by constructing targeted mutations, we found that mutations that increase expression of the outer membrane cobamide transporter BtuB are beneficial during growth under cobamide-limiting conditions. Unexpectedly, we also found that overexpression of the cobamide adenosyltransferase BtuR confers a specific growth advantage in pCbl. Characterization of the latter phenotype revealed that BtuR and adenosylated cobamides contribute to optimal MetH-dependent growth. Together, these findings improve our understanding of how bacteria expand their cobamide-dependent metabolic potential.

**Importance:** In nature, bacteria commonly experience fluctuations in the availability of required nutrients. Thus, their environment often contains nutrients that are insufficient in quantity or that function poorly in their metabolism. Cobamides, the vitamin B_12_ family of cofactors, are ideal for investigating the influence of nutrient quality on bacterial growth. We performed a laboratory evolution experiment in *E. coli* with a less-preferred cobamide to examine whether and how bacteria can improve their growth with less ideal nutrients. We found that overexpression of genes for cobamide uptake and modification are genetic adaptations that improve growth under these conditions. Given that cobamides are key shared metabolites in microbial communities, our results reveal insights into bacterial interactions and competition for nutrients.

## Introduction

Cobamides, the vitamin B_12_ family of metabolites, are used by most bacteria as cofactors for diverse metabolic processes including carbon metabolism, synthesis of methionine and deoxyribonucleotides, and natural product biosynthesis (1). They are produced exclusively by prokaryotes, though most bacteria that use cobamides are incapable of *de novo* synthesis (2, 3). Cobamides are modified tetrapyrroles (corrinoids) with a central cobalt ion that can coordinate to variable upper and lower axial ligands (2). The upper (β) ligand varies depending on catalytic function: a 5ʹ-deoxyadenosyl group is used for radical-based molecular rearrangements, while a methyl group is used for methyltransfer reactions (Fig. 1A). While B_12_ (cobalamin, Cbl) (Fig. 1A) is the best-studied cobamide due to its importance in human health (4), nearly 20 other cobamides with structural variability in the lower (α) axial ligand have been described (5–7). Different cobamides have distinct effects on microbial growth because cobamide-dependent growth and enzyme function are differentially impacted by lower ligand structure (6, 8–15). Diverse assortments of cobamides have been found in microbial communities from host-associated and environmental sources, and variability in cobamide abundances has been observed even in samples derived from similar sources (5, 16, 17). Recent studies have shown that microbial community composition can be significantly altered by addition of certain cobamides, suggesting cobamides can influence microbiomes (18–22). Because microbes can be exposed to different cobamides as their environments shift, they may encounter cobamides that function at varying levels of effectiveness for their metabolism. Given that cobamide structure and availability impact bacterial fitness and community structure, it is important to understand how bacteria are genetically wired to deal with different cobamides.

**Figure 1.**
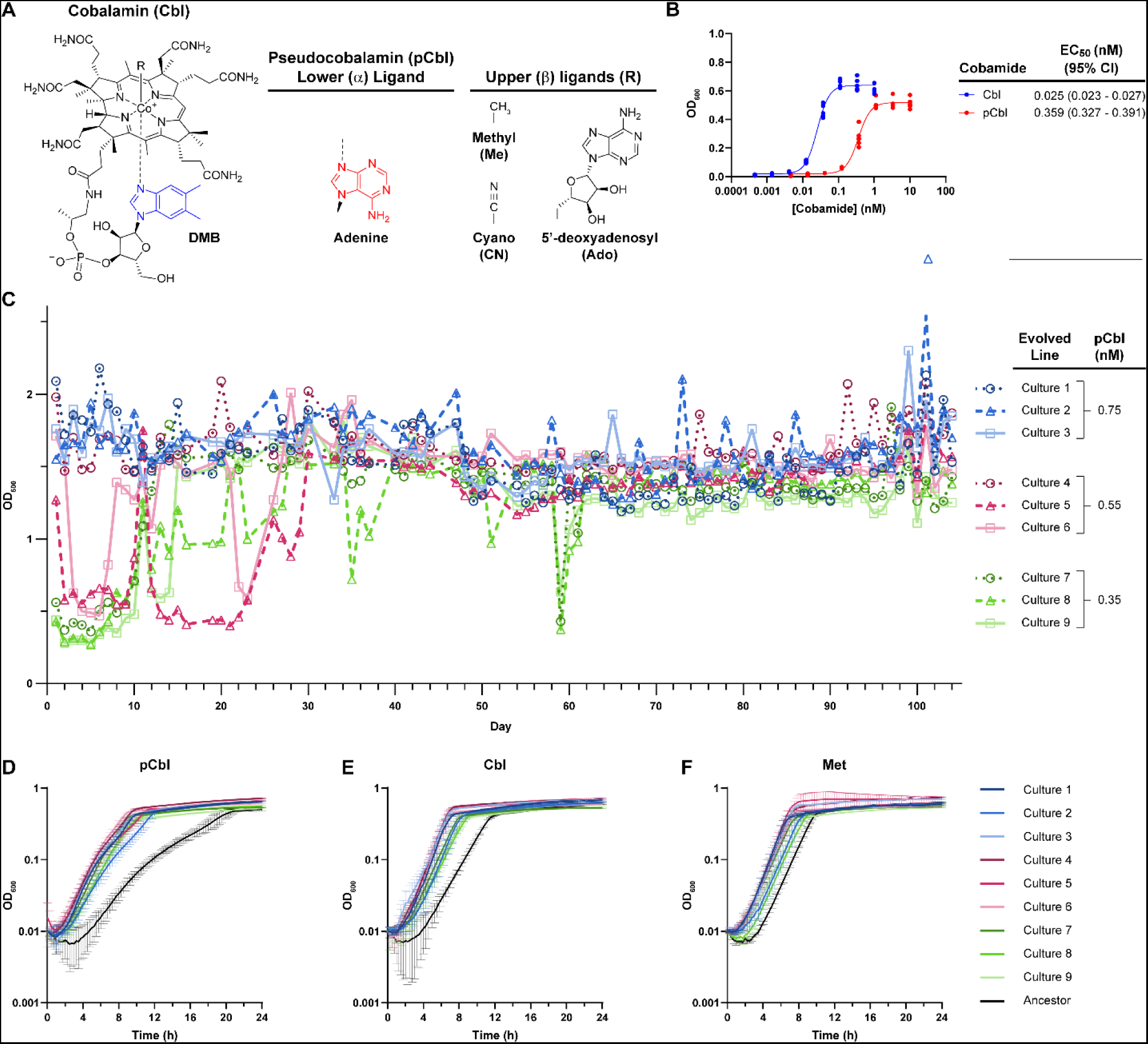
Laboratory evolution of *E. coli* improves its growth with pCbl. A) Structure of cobalamin (Cbl; B_12_) with its lower ligand 5,6-dimethylbenzimidazole (DMB) in blue. Pseudocobalamin (pCbl) contains adenine (red) as its lower ligand. Cobamide upper ligands characterized in this study are shown. B) Dose-response curves of *E. coli* Δ*metE* grown in the absence of methionine with various concentrations of Cbl or pCbl. OD_600_ was recorded after 22 hours. EC_50_ values and 95% confidence intervals of six biological replicates for each cobamide are shown. C) Growth of *E. coli* Δ*metE* cultures during laboratory evolution. Three biological replicate cultures of *E. coli* Δ*metE* were passaged daily in M9 medium supplemented with 0.75, 0.55, or 0.35 nM pCbl for 104 days. OD_600_ was measured prior to diluting 1:100 into fresh medium every 24 hours. D-F) Growth curves of evolved populations (Day 104) and the ancestral Δ*metE* strain in media with 0.35 nM pCbl (D), 0.35 nM Cbl (E), or 0.1 mg/ml Met (F). The average of three biological replicates is shown for panels D-F; error bars represent standard deviation.

Many organisms have evolved strategies to cope with the absence of preferred cobamides. Certain bacteria, archaea, and algae carry out cobamide remodeling, whereby non-preferred cobamides are converted into forms that can be used by their cobamide-dependent enzymes (9, 23–27). In addition, many bacteria encode cobamide-independent alternative enzymes or pathways, circumventing the need for cobamides for certain processes (3, 28). For example, cobamide-independent methionine synthase (MetE) and ribonucleotide reductases are each found in over half of bacterial genomes, including in approximately 80 and 40% of genomes that encode cobamide-dependent counterparts to these enzymes, respectively (3). Bacteria can also tailor their genetic response to the cobamides they prefer via selectivity in riboswitches, noncoding RNA elements in the 5ʹ untranslated region (UTR) of mRNA that, upon binding to specific cobamides, typically downregulate expression of cobamide biosynthesis enzymes, transporters, and cobamide-independent enzymes (29–32).

Here, we carried out a laboratory evolution experiment in *Escherichia coli* to investigate whether there are additional genetic strategies microbes may employ to improve their use of less-preferred cobamides. Due to its short generation time and genetic tractability, *E. coli* is used extensively as a model system to address fundamental biological questions. Although *E. coli* does not require exogenous cobamides in the absence of methionine because it contains *metE*, a Δ*metE* mutant relies on the cobamide-dependent methionine synthase MetH. Adeninylcobamide (pseudocobalamin, pCbl) (Fig. 1A) is a cobamide present in diverse environments, such as the human gut (16), but we find it is used less effectively than Cbl by *E. coli*. In our evolution experiment, we found that, indeed, an *E. coli ΔmetE* mutant can improve its growth with pCbl via several genetic strategies. Different sets of mutations were found in evolved lines provided with different pCbl concentrations, but a common strategy that emerged was increasing the expression of the outer membrane corrinoid transporter BtuB by 300-fold, which provided a competitive advantage in limiting concentrations of cobamides. We additionally found that evolved lines and engineered strains that overexpress the corrinoid adenosyltransferase BtuR are better adapted for growth on pCbl. As a result, this study has revealed a previously unknown role for BtuR in MetH-dependent growth.

## Results

### Laboratory evolution of E. coli improves use of pCbl during MetH-dependent growth

The *E. coli* MG1655 *ΔmetE* strain, which requires MetH activity for growth in minimal medium lacking methionine, prefers Cbl over pCbl, as revealed by comparing growth with the two cobamides (Fig. 1B). Specifically, the concentration of pCbl necessary for half-maximal growth (EC_50_) of this strain is over 10-fold higher than for Cbl, and the maximal growth yield (OD_600_) is lower with pCbl (Fig. 1B). We therefore performed a laboratory evolution experiment with pCbl to determine whether *E. coli* can improve its use of this less-preferred cobamide. Nine independent cultures of the *E. coli* Δ*metE* strain were passaged daily for 104 days for a total of approximately 700 generations in M9 minimal medium containing either 0.75, 0.55, or 0.35 nM pCbl (Fig. 1C). These concentrations encompassed nearly-saturating to limiting growth of the ancestral strain (Fig. 1B). Five of the nine cultures reached a maximal OD_600_ below 0.6 during some or all of the first 10 days, but exceeded an OD_600_ of 1.2 for nearly all passages after day 30, indicating they had adapted to the limiting pCbl conditions (Fig. 1C). When compared to the ancestor, all nine populations showed improved growth with 0.35 nM pCbl (Fig. 1D). The nine populations also showed improved growth with Cbl (Fig. 1E), suggesting that they had evolved better use of cobamides in general. Growth with Met was modestly improved in the evolved populations, indicating they had adapted to other features of the growth medium (Fig. 1F).

### Mutants in one evolved population have a growth advantage specifically with pCbl

We noticed that, when plated on LB agar, which contains methionine, some of the colonies from a passaged culture containing 0.35 nM pCbl (Culture 8) were distinctly smaller than the others (Fig. 2A). These small-colony variants first appeared on day 28, and they made up nearly the whole population on day 65 before being almost entirely lost by day 84 (Fig. 2B). Small-colony variants can arise during laboratory evolution experiments and often are not related to the phenotype being studied, as is the case in this study (see below) (33). Nonetheless, we took advantage of this distinct morphology by using it as a convenient markerless phenotype for further characterizing the pCbl-specific growth advantage of these strains.

**Figure 2.**
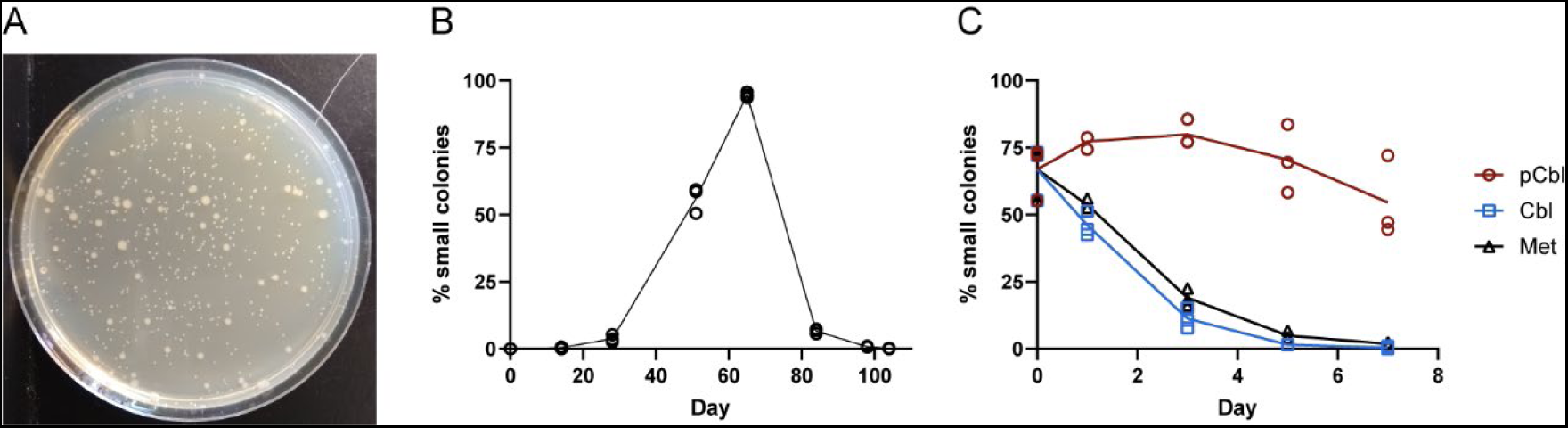
Small colony variants emerge during evolution of Culture 8. A) Plating of a population of Culture 8 on LB agar shows the regular and small colony phenotypes. B) The percentage of total colonies with small size was determined for archived populations of Culture 8. The Δ*metE* ancestor was used for the zero timepoint. C) The percentage of small colonies is plotted for the day 65 archived population of Culture 8 grown with 0.35 nM pCbl, 0.35 nM Cbl, or 0.1 mg/ml Met over seven days with daily passaging. Lines connect the means of three biological replicates.

Growth experiments on the archived culture from day 65 showed that the small-colony variants persisted following one week of daily passaging in the presence of pCbl, whereas their abundance decreased to zero in media containing Cbl or Met (Fig. 2C). The percentage of small colonies in the cultures supplemented with pCbl started to decrease on day 5 of the experiment (Fig. 2C), suggesting that the same population dynamics may have been occurring that were seen during the laboratory evolution (Fig. 2B).

We isolated colonies with different sizes from the day 65 population and individually competed three “small” isolates (S2, S3, and S4) against two “regular” isolates (R1 and R3), as well as the ancestral Δ*metE* strain, in media containing either pCbl, Cbl, or Met. All three small isolates had similar phenotypes. When co-cultured, the small isolates outcompeted the ancestor in the presence of cobamides, taking over the entire population after a single passage with pCbl and after three passages with Cbl (Fig. 3A, B, Fig. S1 A, B, D, E). In media with methionine, however, the ancestral strain outcompeted the evolved isolates (Fig. 3C, Fig. S1 C, F). When the small isolates were competed against the two regular isolates, the small isolates were outcompeted when grown with Cbl and Met, but we observed contrasting phenotypes with pCbl (Fig. 3 A-C, Fig. S1). The small isolates had a competitive advantage over regular isolate R1 but were outcompeted by R3 (Fig. 3A, Fig. S1 A, D).

**Figure 3.**
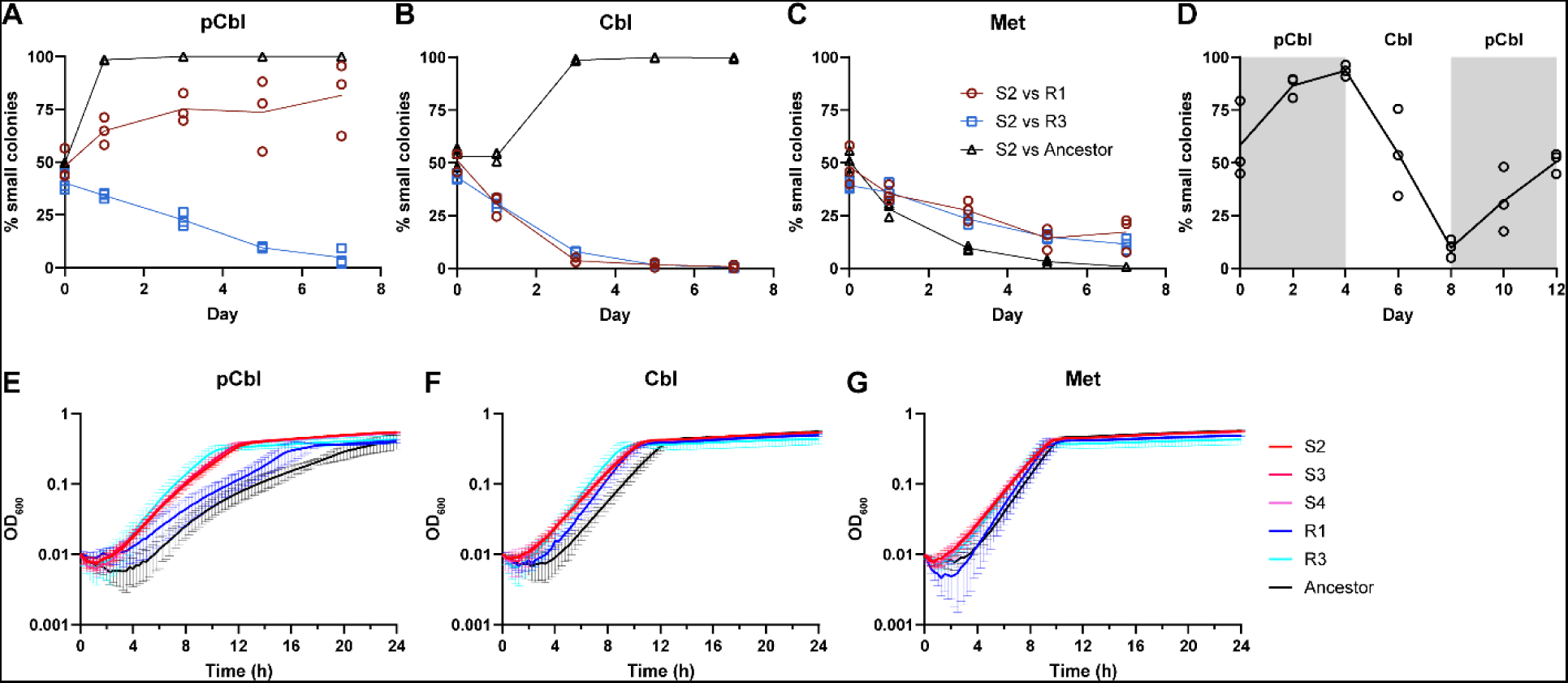
Growth characteristics of isolates S2, R1, and R3 from Culture 8. A-C) Isolate S2 was competed against isolates R1 and R3 and the ancestor strain for seven days with daily passaging in medium containing 0.35 nM pCbl (A), 0.35 nM Cbl (B), or 0.1 mg/ml Met (C). Cultures were diluted and plated on the indicated days to quantify the fraction of small colonies, corresponding to S2 strain abundance. D) Isolates S2 and R1 were competed for 12 days with daily passaging in medium containing either 0.35 nM pCbl (shaded) or 0.35 nM Cbl (unshaded), and the fraction of small colonies (S2) was calculated as in panels A-C. Lines in A-D connect the means of three biological replicates. E-G) Growth curves of isolates S2, S3, S4, R1, R3 and the ancestor strain in media containing 0.35 nM pCbl (E), 0.35 nM Cbl (F), or 0.1 mg/ml Met (G). The average of three biological replicates is shown; error bars represent standard deviation.

The competitive advantage of the small isolates with pCbl was further confirmed by co-culturing isolates S2 and R1 in medium supplemented alternately with pCbl and Cbl. When passaged with pCbl for four days, the proportion of S2 increased to over 90%, but after switching to the Cbl-containing medium, the proportion of S2 decreased to less than 10% after four days. A subsequent return to pCbl resulted in an increase in S2 (Fig. 3D).

The phenotypes of the small and regular isolates that we observed in competition were consistent with their growth characteristics in pure culture (Fig. 3E-G). Isolates S2, S3, and S4, which outcompeted the ancestor in media with pCbl and Cbl but were outcompeted with Met, grew faster than the ancestor in the presence of cobamides while showing a small growth improvement with Met (Fig. 3E-G). Only isolate R1, which competed poorly in the presence of pCbl, had less improved growth than the other isolates, particularly in the medium containing pCbl (Fig. 3E). Isolate R3, which outcompeted the small isolates in all three media conditions, grew similarly to the small isolates in each medium (Fig. 3E-G). Taken together, these results suggest that all of the isolates have acquired one or more mutations that confer a growth advantage with cobamides. Further, based on the growth phenotypes of strains S2, S3, S4 and R3 with pCbl, these strains likely have one or more mutations that confer a specific advantage with pCbl.

### Evolved isolates have mutations that increase expression of cobamide-related genes

To identify the mutations acquired during the evolution experiment, we performed whole genome sequencing on the isolates from Culture 8. Each isolate has a unique set of mutations, which range from 8 to 14 single nucleotide polymorphisms (SNPs), insertions and deletions (InDels), or structural variants (SVs) (Fig. 4A; Table S1). This includes mutations in known cobamide-related genes in all five isolates. They each have the same two SNPs in the promoter and ribosome binding site (RBS) of the *btuB-murI* operon, which encodes the outer membrane corrinoid transporter BtuB and the glutamate racemase MurI (Fig. 4A). We predict that these mutations result in increased expression of the operon (see Fig. S2A for details). RT-qPCR on the ancestor strain and the evolved isolates confirmed that the operon is overexpressed in the evolved isolates. *btuB-murI* transcript levels in the evolved isolates were approximately 300-fold higher than in the ancestor (Fig. 4B).

**Figure 4.**
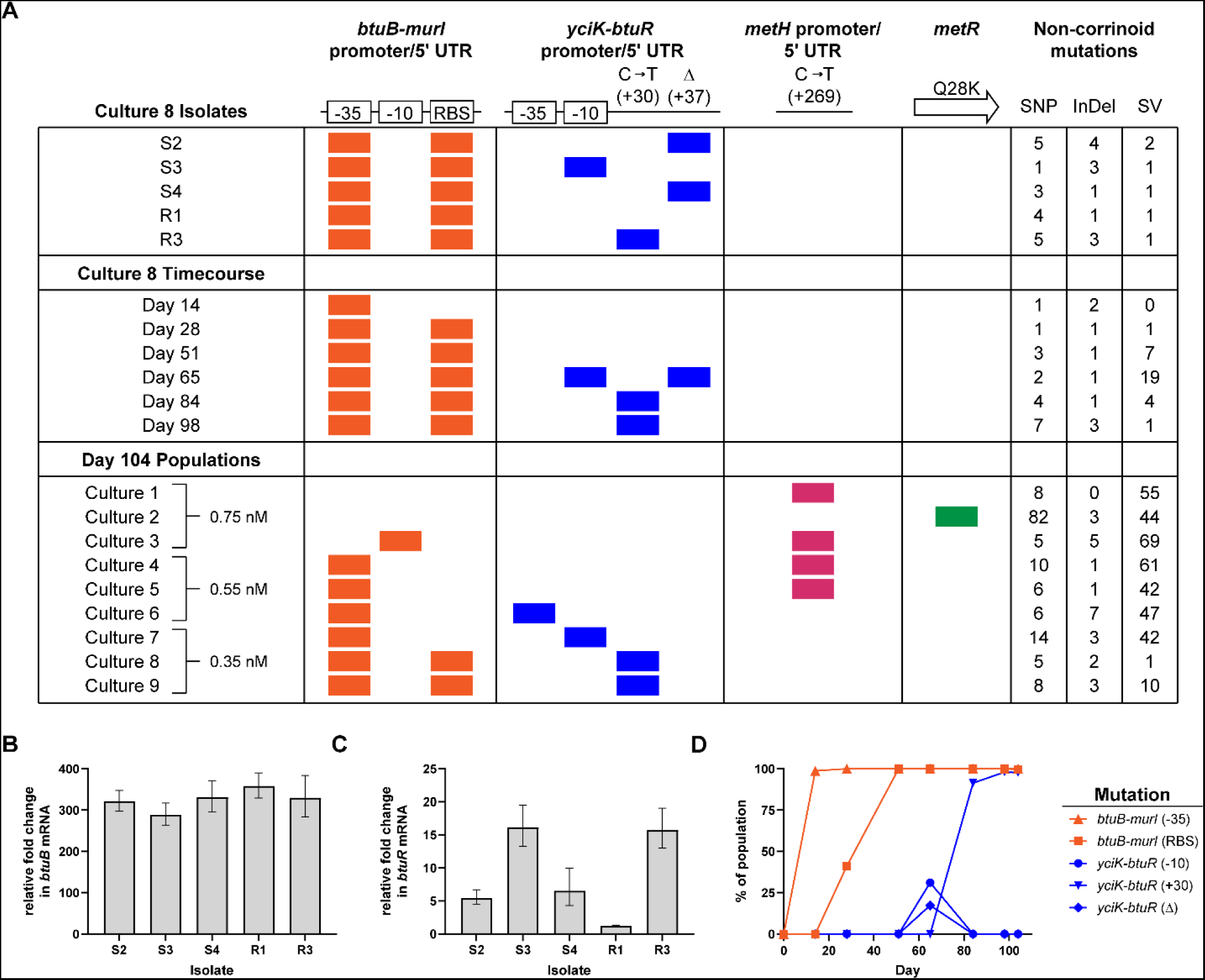
Corrinoid-related genes are mutated and upregulated during evolution. A) Colored boxes show the presence of the indicated mutations affecting corrinoid-related genes in isolates and archived timepoints from Culture 8, and in the endpoint (Day 104) populations of all evolved lines. The promoter for the *yciK-btuR* operon has not been experimentally characterized and was predicted by PromoterHunter (67). The specific changes in the promoters and 5ʹ UTRs of *btuB-murI* and *yciK-btuR* are shown in Figure S2. *yciK-btuR* is on the minus strand of the chromosome and the +30 change (C to T) is the reverse complement of the mutation in the 5ʹ UTR. The concentration of pCbl present during the evolution of each line (Cultures 1-9) is indicated. The numbers of SNPs, InDels, and SVs affecting non-corrinoid-related genes in each sequenced isolate or population are shown (see Table S1 for list of genes and mutations). B-C) *btuB* (B) and *btuR* (C) transcript levels in evolved isolates relative to Δ*metE* ancestor from cultures containing 1 nM pCbl as determined by RT-qPCR. Mean and error were calculated from three biological replicates. D) Frequency of the *btuB-murI* and *yciK-btuR* mutations within the Culture 8 population during the evolution experiment.

In addition to the mutations in *btuB-murI*, all of the isolates except R1 have a mutation in the predicted promoter or 5ʹ UTR of the *yciK-btuR* operon (Fig. 4A, Fig. S2B). *yciK* is an uncharacterized gene annotated as a putative oxidoreductase, and *btuR* encodes an adenosyltransferase that installs a 5ʹ-deoxyadenosyl group as the β ligand of corrinoids. RT-qPCR analysis of the evolved isolates showed that isolates with *btuR* mutations (S2, S3, S4, and R3) had 5- to 15-fold higher levels of *yciK-btuR* mRNA compared to the ancestor, while mRNA levels similar to the ancestor were observed in isolate R1, which does not have a mutation in *yciK-btuR* Fig. 4B). These results suggest that growth of *E. coli* with limiting amounts of pCbl can be improved by increasing the expression of these two operons.

Sequencing of the archived populations of Culture 8 enabled us to follow the emergence of these mutations during the evolution experiment (Fig. 4A, D; Table S1). Both *btuB-murI* mutations arose early in the timecourse and were retained within the entire population by day 51 (Fig. 4D). Notably, mutations in the −35 element and RBS were present by days 14 and 28, respectively, coinciding with increases in the OD_600_ of the culture (Fig. 1C).

The two *yciK-btuR* mutations found in the small isolates were first detected on day 65, in 31 and 17% of the population, respectively (Fig. 4D). At all of the following timepoints, however, only the C to T mutation found in isolate R3 was detected in the population. Given that isolate R3 outcompeted isolates S2, S3 and S4 in pCbl (Fig. 3A, Fig. S1), it is likely that descendants of R3 became dominant in the population after day 65. An *rpsK* mutant was observed to have a small colony phenotype in a previous study in *E. coli* (34), and we found here that the frequency of an *rpsK* mutation correlated with the prevalence of small colonies in the population (Table S1).

### *E. coli* adapts differently with limiting versus near-saturating pCbl

The endpoint (Day 104) archives of the nine evolved cultures were also sequenced. The populations have several of the cobamide-associated mutations found in the Culture 8 isolates and earlier timepoints, as well as mutations in nearly 400 additional genes, including 75 genes found mutated in more than one sample (Fig. 4A; Table S1). In each population, at least one known cobamide-related gene is mutated, with different genes affected depending on the concentration of pCbl present during the evolution (Fig. 4A). The two other populations passaged with 0.35 nM pCbl developed the same mutations upstream of *btuB-murI* and *yciK-btuR* as those found in Culture 8 (Fig. 4A, Cultures 7 and 9). In contrast, only a minor fraction (14%) of one population passaged with 0.75 nM pCbl has a mutation upstream of *btuB-murI*, and one population passaged with 0.55 nM pCbl has a mutation upstream of *yciK-btuR* (Cultures 3 and 6, respectively; Fig. 4A; Table S1), though all three populations grown with 0.55 nM pCbl have the same *btuB-murI* promoter mutation found in the isolates.

All of the evolved populations lacking mutations in *yciK-btuR* have a mutation upstream of *metH* or in the coding sequence of *metR*, a transcriptional activator of *metH* (35), in 50% or more of the population (Culture 1-5; Fig. 4A; Table S1). It is unclear how these mutations affect *metH* expression; the *metH* mutation is not located in the promoter, RBS, or MetR binding site (36), while the *metR* mutation is located in its DNA-binding domain (37). Taken together, these results suggest that increasing cobamide uptake and adenosylation are effective strategies for improving growth with limiting to moderate pCbl concentrations, while changing expression of *metH* facilitates adaptation at higher concentrations of pCbl.

### Overexpression of the corrinoid uptake gene btuB is advantageous at limiting cobamide concentrations

Having found that the operons containing *btuB* and *btuR* are upregulated in the evolved isolates, we next examined how overexpression of each gene separately affects cobamide-dependent growth. We first investigated the influence of *btuB* alone, since the glutamate racemase MurI, encoded in the same operon, has no known function in cobamide metabolism (38). We constructed a strain with a Δ*metE* mutation that overexpresses *btuB* by inserting a second copy of the gene into the chromosome, with its promoter containing the G to T mutation found in the −35 element of the evolved isolates. This strain was competed against a Δ*metE* control strain, with each strain expressing either CFP or YFP to monitor their abundances in co-culture. We found that overexpression of *btuB* conferred a competitive advantage in media containing 1 nM pCbl, but not 1 nM Cbl or Met (Fig. 5 A-C). However, competing the strains in varying cobamide concentrations showed that *btuB* overexpression is beneficial with both pCbl and Cbl, but only at concentrations at which the cobamide is limiting, namely 1 nM and less for pCbl, and under 0.25 nM for Cbl (Fig. 5D, E). Thus, the *btuB* mutations that arose during passaging with limiting pCbl presumably improved *E. coli*’s ability to import cobamides to support MetH-dependent growth.

**Figure 5.**
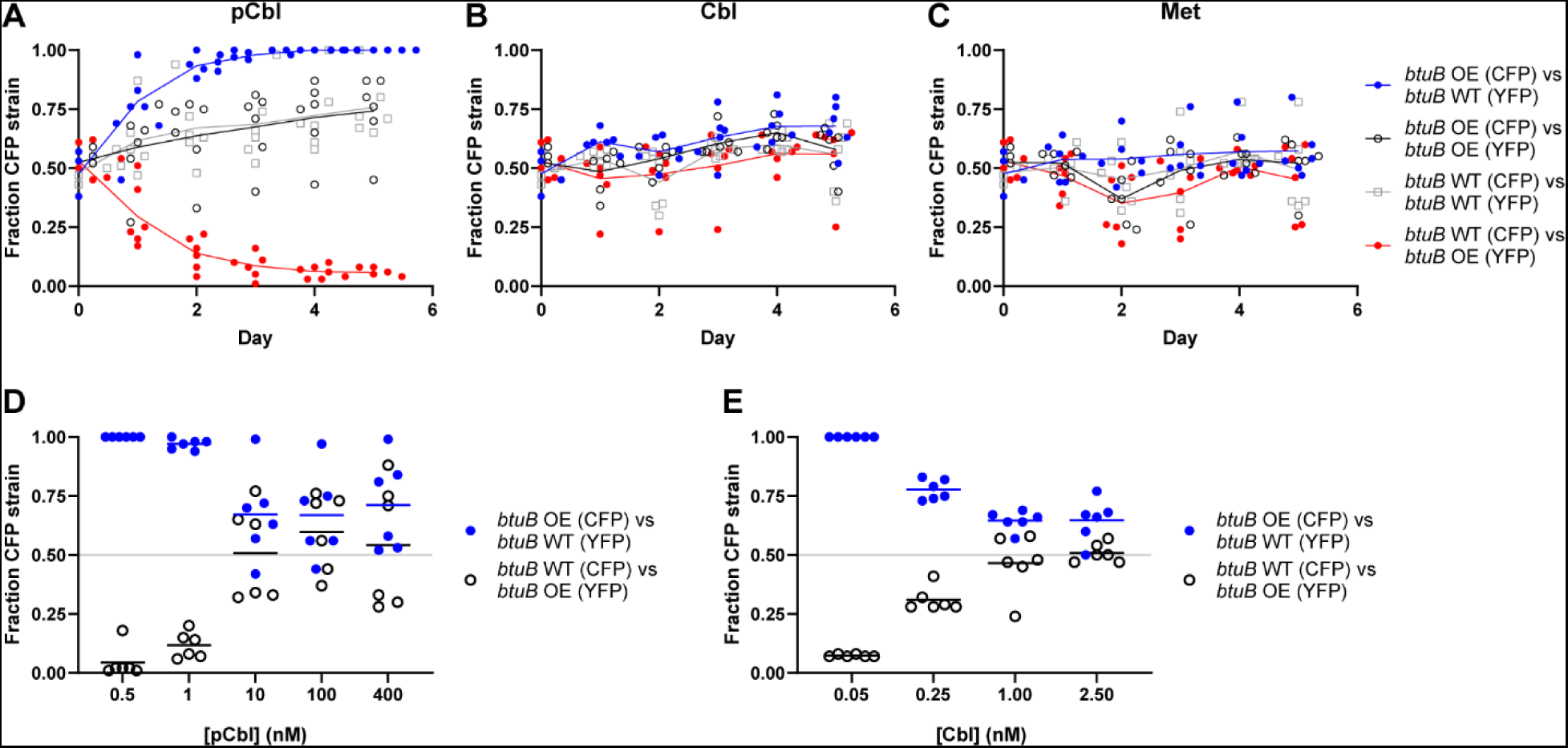
Overexpression of *btuB* confers a competitive advantage at limiting cobamide concentrations. A-C) *E. coli* Δ*metE* strains overexpressing *btuB* (OE) or producing native levels of *btuB* (WT), each expressing either CFP or YFP, were competed in co-culture for five days with daily passaging in medium containing either 1 nM pCbl (A), 1 nM Cbl (B), or 0.1 mg/ml Met (C). The fraction of the CFP-expressing strain in each co-culture is plotted. Control co-cultures containing CFP- and YFP-expressing strains in the same genetic background (black and gray) were included to rule out a growth disadvantage caused by either fluorescent protein. D-E) The CFP- and YFP-expressing strains that overexpress *btuB* (OE) or produce native levels of *btuB* (WT) were competed in co-culture at different concentrations of pCbl (D) or Cbl (E). Fluorescence was measured on day 3 following daily passaging. Lines represent the means of six biological replicates.

### The corrinoid adenosyltransferase gene btuR is required for optimal MetH-dependent growth

Next, we assessed whether increasing *btuR* expression impacts MetH-dependent growth by overexpressing *btuR* on a plasmid. We found that, similar to the results with *btuB*, a strain overexpressing *btuR* outcompeted a strain with wild type *btuR* levels when grown with pCbl, but not with Cbl or Met (Fig. 6 A-C). Though it is in an operon with *yciK*, *btuR* alone was responsible for this phenotype, as overexpression of *yciK* did not confer a growth advantage with pCbl and co-expression of *yciK* with *btuR* did not influence the effect of overexpression of *btuR* alone (Fig. S3). However, unlike *btuB*, overexpression of *btuR* conferred a competitive growth advantage even at concentrations of up to 400 nM pCbl, but failed to confer an advantage at any concentration of Cbl tested (Fig. 6 D, E).

**Figure 6.**
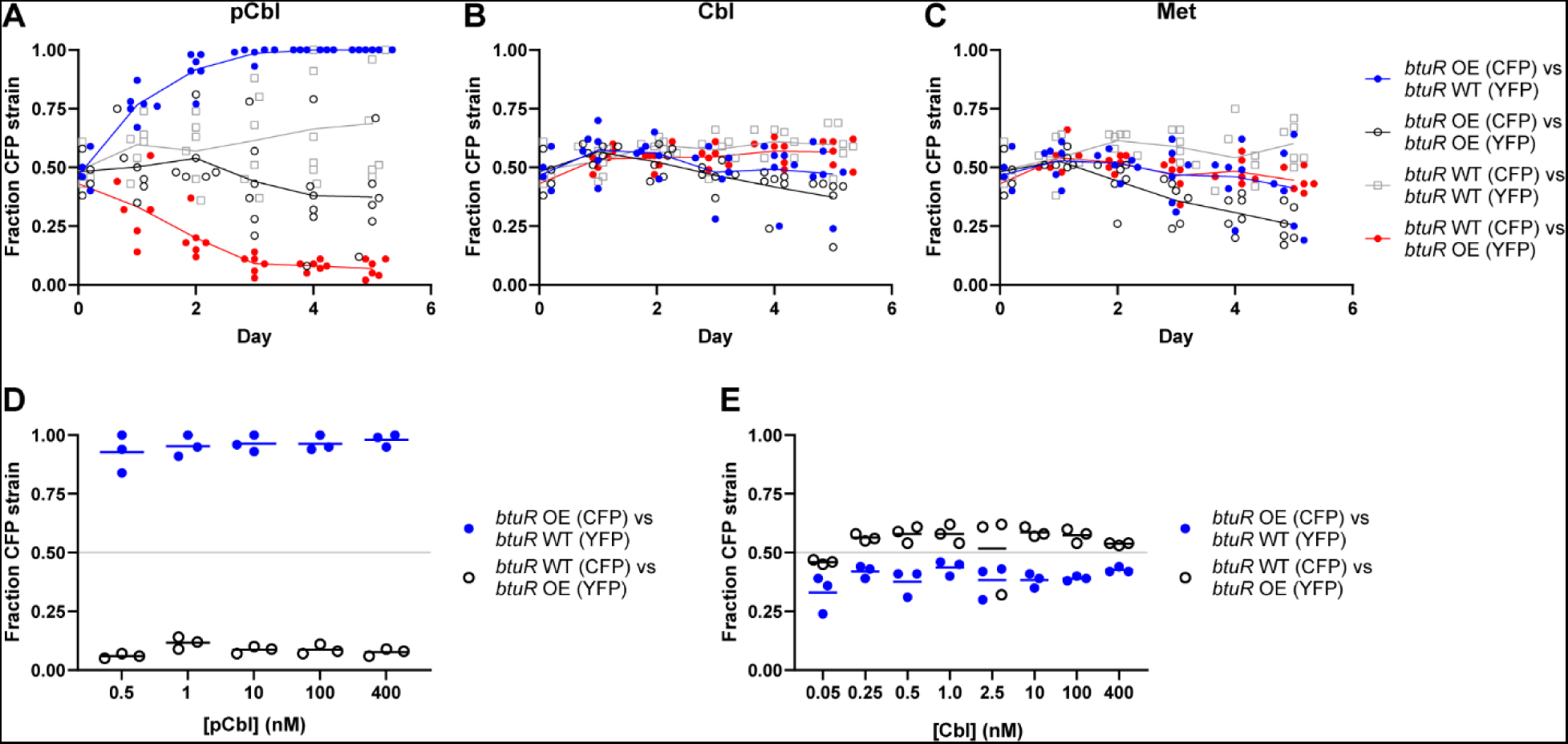
Overexpression of *btuR* confers a competitive advantage only during growth with pCbl. A-C) *E. coli* Δ*metE* strains overexpressing *btuR* (OE) or producing native levels of *btuR* (WT), each expressing CFP or YFP, were competed in co-culture for five days with daily passaging in medium containing either 1 nM pCbl (A), 1 nM Cbl (B), or 0.1 mg/ml Met (C). The fraction of the CFP-expressing strain in each co-culture is plotted. Control co-cultures containing CFP- and YFP-expressing strains in the same genetic background (black and gray) were included to rule out a growth disadvantage caused by either fluorescent protein. D-E) The CFP- and YFP-expressing strains that overexpress *btuR* (OE) or produce native levels of *btuR* (WT) were competed in co-culture at different concentrations of pCbl (D) or Cbl (E). Fluorescence was measured on day 3 following daily passaging. Lines represent the means of six biological replicates.

In the Δ*metE* mutant, cobamides are used by the MetH enzyme to transfer methyl groups from methyltetrahydrofolate to homocysteine by alternately methylating and demethylating the cobamide at the β position. It was therefore puzzling to find that overexpression of BtuR, which adenosylates cobamides at the β position, improves MetH-dependent growth. To further explore the role of BtuR in MetH-dependent growth, we deleted *btuR* and performed growth assays with pCbl or Cbl with either cyano (CN, as in Fig. 1-6) or adenosyl (Ado) β ligands (Fig. 1A). Growth measurements with these cobamides showed that a Δ*btuR* Δ*metE* strain has impaired growth in cyanopseudocobalamin (CNpCbl), with a lower maximum OD_600_ and an EC_50_ over 25-fold higher than the Δ*metE* strain (Fig. 7A). Growth with adenosylpseudocobalamin (AdopCbl) led to a higher maximum OD_600_ and lower EC_50_ of the Δ*btuR* Δ*metE* strain, though growth was still considerably impaired compared to the Δ*metE* strain (Fig. 7A). A similar trend was observed when these strains were cultured with cyanated versus adenosylated forms of Cbl (CNCbl and AdoCbl, respectively), though the growth impairment of the Δ*btuR* Δ*metE* strain was more modest (Fig. 7B). Together, these results confirm that *btuR* influences MetH-dependent growth in *E. coli*.

**Figure 7.**
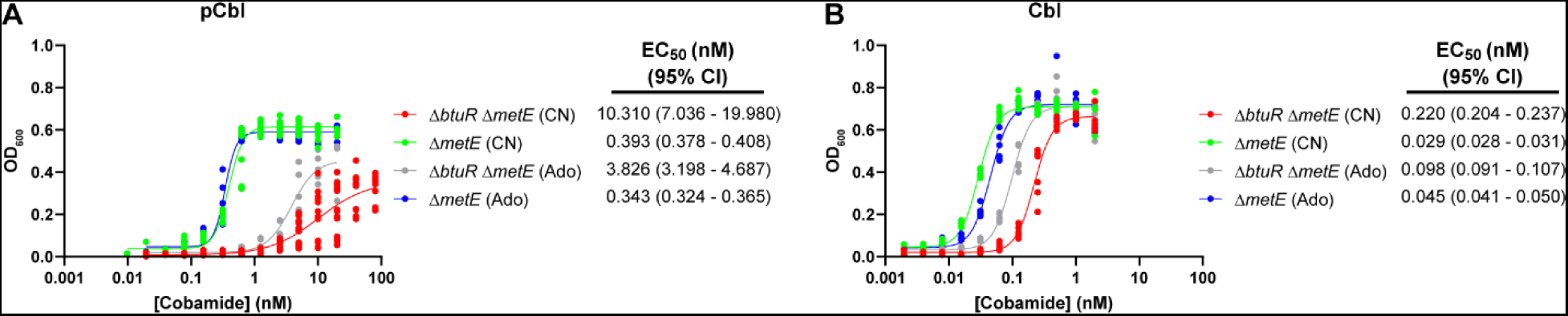
Deletion of *btuR* causes poorer growth with pCbl and Cbl. A-B) Cobamide dose-response curves are shown for *E. coli* Δ*btuR* Δ*metE* and Δ*metE* strains grown with various concentrations of CNpCbl and AdopCbl (A), or CNCbl and AdoCbl (B) with no added methonine. OD_600_ was recorded after 22 hours of growth. EC_50_ values and 95% confidence intervals were calculated from 6-18 biological replicates for each cobamide concentration.

The growth defect of the Δ*btuR* Δ*metE* strain and partial rescue by adenosylated cobamides could be due to differences in cobamide uptake or retention. To examine this possibility, we measured the amount of cobamide in the supernatant and cell pellet fractions in the Δ*btuR* Δ*metE* and Δ*metE* strains cultured with each of the four cobamides tested in Figure 7 (AdopCbl, CNpCbl, AdoCbl, CNCbl) using a quantitative bioassay that detects corrinoids (cobamides and biosynthetic precursors). Strains lacking *btuR* had approximately twofold less AdopCbl and CNpCbl in the cell pellet fraction (Fig. S4A); this result was validated by HPLC analysis of corrinoid extractions (Fig. S4D). No differences in the levels of either AdoCbl or CNCbl was observed between the Δ*btuR* Δ*metE* and Δ*metE* strains, but less AdoCbl was found in the cell pellets than CNCbl (Fig. S3 B, C). The twofold differences in intracellular pCbl levels between the strains may only partially explain the 10- to 25-fold differences in EC_50_ and differences in maximal OD_600_ observed in the dose-response assays (Fig. 7A). Thus, it is possible that the adenosylated forms of cobamides play a previously unrecognized role in promoting MetH activity.

## Discussion

Cobamides are considered key shared nutrients because they function as cofactors for numerous microbial processes but are only produced by a subset of prokaryotes. They have been detected in diverse microbial communities, both environmental and host-associated, and a wide range in cobamide levels has been observed across ecosystems, with some dominated by one or two cobamides, while others contain up to eight different types (16). These differences in cobamide diversity across environments are noteworthy in light of the observation that many bacteria have preferences for particular cobamides and most bacteria rely on cobamides produced by other microbes. This raises the question of how bacteria adapt in the presence of non-preferred cobamides. We addressed this question by using a cobamide-dependent mutant of *E. coli* in a laboratory evolution experiment. We found that *E. coli* is indeed capable of improving its growth with pCbl via genetic changes, and it uses differing strategies depending on the availability of the nutrient. Competition experiments and genetic analyses revealed regulation of corrinoid uptake as a limiting factor in *E. coli* and a previously unappreciated role for the corrinoid adenosyltransferase BtuR in MetH-dependent growth.

We previously showed that the cobamide-dependent enzyme methylmalonyl-CoA mutase (MCM) has different binding affinities for different cobamides, and that these cobamide-binding affinities largely mirror MCM-dependent growth with different cobamides in *Sinorhizobium meliloti* (13). Other studies have shown that MetH orthologs from different organisms display distinct preferences for different cobamides (9, 14, 25). Given that pCbl is less preferred by *E. coli* than Cbl (Fig. 1B), we hypothesized that passaging with pCbl would lead to the accumulation of mutations in the MetH enzyme that improved its ability to bind and use pCbl. Though we observed mutations that presumably impact *metH* expression, no mutations were found in the *metH* coding sequence. Altering expression of corrinoid-related genes was the general outcome of our evolution experiment, suggesting that modifying the regulation of cobamide metabolism may be a more readily accessible mechanism of adaptation than changes to the specificity of the dependent enzyme, particularly in our experimental timeframe. Changes to gene expression are routinely seen in laboratory evolution experiments, including the targeting of global regulators (39, 40).

While mutations in *metH* and its transcriptional activator *metR* were found at the higher concentration of pCbl, mutations upregulating the outer membrane corrinoid transporter BtuB arose primarily when pCbl was limiting, consistent with its role in corrinoid uptake. Indeed, we previously showed that overexpression of the corrinoid transport machinery in *Bacillus subtilis* increases the amount of cobamide imported (32). Although it is not known whether the affinity of BtuB for Cbl versus pCbl differs, structural and biochemical studies with Cbl and a corrinoid intermediate lacking a lower ligand (cobinamide) have shown that the lower ligand is of minimal importance to binding, with BtuB primarily interacting with the corrin moiety (41–43). Instead, greater *btuB* expression, leading to increased pCbl uptake, could compensate for decreased MetH efficiency with pCbl at a later step such as adenosylation by BtuR or utilization by MetH. Alternatively, the evolved mutations upregulating *btuB* may also be a consequence of the selective pressure from competing for limiting pCbl. Cobamide uptake is critical for colonization by the human gut commensal bacterium *Bacteroides thetaiotaomicron* in a mouse model (44), and *B. thetaiotaomicron* and other *Bacteroides* species encode multiple corrinoid transport systems which include high-affinity corrinoid binding proteins absent from *E. coli* (45). These uptake systems are thought to enable *Bacteroides* to outcompete other microbes for corrinoids, allowing for successful gut colonization (46, 47). Our observation that corrinoid uptake and competitiveness in *E. coli* can readily be improved via mutations in the *btuB* promoter or RBS suggests that *E. coli* is not evolved to maximize corrinoid uptake, despite its being a member of the gut microbiota like *B. thetaiotaomicron*. This is notable given that the purinyl cobamides, which include pCbl, were found to be the dominant fecal corrinoids in the majority of human subjects (5). *E. coli* could be under less selective pressure to maximize corrinoid uptake because, unlike *B. thetaiotaomicron*, *E. coli* has the cobamide-independent methionine synthase *metE* as well as *metH*, rendering it less dependent on exogenous cobamides. In addition, BtuB is a phage receptor in *E. coli*, so increased expression of *btuB* may not always be beneficial in natural settings (48).

The BtuR corrinoid adenosyltransferase is responsible for installing a 5ʹ-deoxyadenosine moiety as the β ligand of cobamides to produce adenosylcobamides (49), which are required for the subset of cobamide-dependent enzymes, such as MCM, that carry out radical-based reactions (50). However, no role for adenosylcobamides has been proposed for methyltransferases such as MetH, which use methylcobamides – cobamides with a methyl group as the β ligand – to shuttle methyl groups from a methyl donor to a substrate. Therefore, it was surprising to find that, in MetH-dependent *E. coli*, overexpression of *btuR* provides a competitive advantage during growth with pCbl, deletion of *btuR* impairs growth with both pCbl and Cbl, and supplementation of the Δ*btuR* mutant with adenosylcobamides only partially rescues the growth phenotype. These results suggest that adenosylcobamides, and perhaps the BtuR protein itself, could have previously unknown roles in MetH function. Some cobamide-dependent enzymes such as MCM require a corrinoid adenosyltransferase and other accessory proteins to load the cobamide cofactor into the enzyme (51–53). It is possible that BtuR fulfills such a role for MetH in *E. coli*, particularly for cobamides that function poorly as a cofactor for MetH. Alternatively, adenosylcobamides and/or BtuR could facilitate SAM-dependent cobamide reactivation, a step required approximately every 2,000 turnovers for Cbl following spontaneous cofactor oxidation (54–56). Until recently, studies of *E. coli* MetH have been unable to address the cofactor loading step because the enzyme is stable only when pre-loaded with Cbl during purification. Future *in vitro* studies with a newly identified MetH homolog that is stable in its apo form will facilitate analysis of this phase of the reaction (57). Because pCbl functions more poorly than Cbl in *E. coli* MetH-dependent growth, our evolution experiment may have fortuitously uncovered a role for adenosylated cobamides in corrinoid-dependent physiology. Future work will be aimed at understanding the molecular mechanisms underlying these observations.

## Material and Methods

### Media and growth conditions

*E. coli* MG1655 Δ*metE* evolution was performed at 37°C with aeration in M9 glycerol (0.2%) minimal medium (58) with the indicated concentrations of pCbl. 20 ml cultures were grown in glass test tubes with 0.2 ml transferred into fresh media every 24 hours. A sample of each population was archived on days 14, 28, 51, 65, 84, 98, and 104 in 25% glycerol and stored at −80°C. Before the start of the evolution experiment, the three replicate cultures were passaged daily for 16 days with a saturating level of pCbl (5 or 2.5 nM) while the appropriate concentrations for the evolution experiment were being determined.

M9 medium was supplemented with L-methionine (Met) at 0.1 mg/ml unless indicated. LB agar was used as solid medium. For experiments with the *E. coli* Δ*metE* ancestor or evolved isolates from Culture 8, M9 medium was inoculated with individual colonies grown on LB agar. Pre-culturing of populations and strains in M9 medium was performed at 37°C with aeration.

### Strain construction

All strains used for evolution and mutant validation are derivatives of wild type K12 strain MG1655. *E. coli* strains were cultured at 37°C with aeration in LB medium during strain construction. Media were supplemented with antibiotics at the following concentrations when necessary: kanamycin, 25 mg/liter (pKIKO, pETmini); carbenicillin, 100 mg/liter (pCP20); chloramphenicol, 10-20 mg/liter (pACYCDuet-1). pKIKO*arsB*Km plasmids were propagated in *E. coli* DH5α containing λ*pir*.

The Δ*metE*::kan^R^ and Δ*btuR*::kan^R^ mutations from the Keio collection (59) were introduced by P1 transduction into *E. coli* strain MG1655 (59, 60). The kanamycin resistance cassette was removed from Δ*metE*::kan^R^ by introducing the plasmid pCP20 as described, leaving the FRT site in place of the *metE* coding sequence (61).

An *E. coli* strain overexpressing *btuB* was created by integrating an additional copy of *btuB* at the *arsB* (arsenite transporter) locus using the KIKO system as described (62). pKIKO*arsB*Km was a gift from Lars Nielsen & Claudia Vickers (Addgene plasmid # 46766; http://n2t.net/addgene:46766; RRID:Addgene_46766). *E. coli btuB* with its promoter and riboswitch was cloned into pKIKO*arsB*Km, with the promoter containing the −35 element mutation (T<u>T</u>GACA) found in evolved populations and isolates. *btuB* also contained a synonymous mutation in codon 581 encoding a valine (GTT to GTA). The PCR-based method was used to integrate this construct at the *arsB* locus. The kanamycin resistance cassette alone from pKIKO*arsB*Km was integrated at the *arsB* locus to create a control strain. Both constructs were first integrated into MG1655 before being transduced via P1 into the Δ*metE* strain. Finally, the kanamycin resistance cassette was removed using pCP20. Strains were confirmed by PCR and Sanger sequencing. Plasmids pSG013 and pSG015, which contain mCerulean and mCitrine (abbreviated CFP and YFP in text), respectively, were each transformed into the two strains to enable tracking based on fluorescence (63).

The *btuR* and *yciK* genes were overexpressed in a pACYCDuet-1 plasmid in which the T7 promoters were replaced with a *lac* promoter and operator (pACYCDuet-1-pLac). This enabled repression of gene expression in the presence of glucose (0.02% in LB, 0.2% in M9) and expression in the absence of glucose due to the leakiness of the *lac* promoter. *E. coli btuR*, *yciK*, and the *yciK-btuR* operon were each cloned downstream of the *lac* promoter in pACYCDuet-1-pLac. mCerulean and mCitrine genes from pSG013 and pSG015 (with J23100 promoter and B0034 RBS) were inserted between the chloramphenicol resistance cassette and p15A origin in each of these plasmid constructs to enable tracking of strains by fluorescence measurements (63).

### Cobamide reagents

CNCbl and AdoCbl were purchased from MilliporeSigma. CNpCbl was extracted from *Propionibacterium acidi-propionici* strain DSM 20273 and purified as described (64, 65). AdopCbl was chemically synthesized from CNpCbl and purified as described (13). Cobamides were quantified spectrophotometrically (13, 26). Cbl and pCbl were used in their cyano forms (CNCbl and CNpCbl) unless otherwise indicated.

### Growth assays and competition experiments

To quantify the percentage of small colonies present during the evolution of Culture 8, archived populations were cultured overnight in M9 glycerol medium supplemented with 0.35 nM pCbl, diluted, and plated on LB agar.

Growth assays and competition experiments were performed in 200 μl cultures in 96-well plates (Corning, 3598). For growth curves, populations or isolates were pre-cultured in M9 glycerol supplemented with 0.35 nM pCbl, while cultures for cobamide dose-response assays were supplemented with Met. Cells from saturated cultures were collected by centrifugation, resuspended in M9 glycerol, and OD_600_ was measured. Each culture was then inoculated at a starting OD_600_ of 0.01 in M9 glycerol medium with the indicated supplement. 96-well plates were sealed with Breathe-Easy (Diversified Biotech). Growth assays were performed in a BioTek Synergy 2 microplate reader with shaking at medium speed at 37°C and OD_600_ recorded every 15 min for 24 hours. OD_600_ for cobamide dose-response assays was measured with the BioTek Synergy 2 microplate reader following 22 h growth at 37°C with shaking in either the plate reader or a heated benchtop microplate shaker (1,200 rpm, Southwest Science). Preparation of cultures containing adenosylcobamides was done under red light and the plates were incubated in the dark. EC_50_ values were calculated using Graphpad Prism (Dose-response – Stimulation; [Agonist] vs. response – Variable slope (four parameters)).

For competition experiments involving evolved populations or strains, cells were pre-cultured in M9 glycerol medium supplemented with Met. Cells were pelleted, washed twice with 0.85% saline, and resuspended in M9 glycerol medium, with the exception of the experiments shown in Fig. 2C and 3D, in which the cells were pelleted and resuspended in M9 glycerol medium without washing. OD_600_ was measured and the population or an equal ratio of two strains was inoculated at a starting OD_600_ of 0.01 in 200 μl M9 glycerol medium containing the indicated supplement. A dilution of the culture was plated on LB agar to establish the percentage of small colonies at time 0. The plate was sealed and incubated at 37°C in a benchtop microplate shaker at 1,200 rpm. 2 μl of each culture was transferred into 198 μl fresh medium every 24 h. On the indicated days, dilutions from the cultures were plated on LB agar to determine the percentage of small colonies in the population.

Competition experiments involving *btuB*-overexpression strains were tracked by fluorescence (63). Strains were pre-cultured in M9 glycerol medium supplemented with Met. Cells were pelleted, washed twice with 0.85% saline, and resuspended in M9 glycerol medium. OD_600_ was measured and each sample was adjusted to an OD_600_ of 0.25. Co-cultures were prepared by mixing an equal volume of each strain. 100 μl of each co-culture was transferred to a 96-well glass bottom plate (P96-1.5P, Cellvis) and cyan and yellow fluorescence were measured on a multiwell plate reader (Tecan Spark) as described (63). Separately, 8 μl of each mono- and co-culture were added to 192 μl of M9 glycerol medium (starting OD_600_ of 0.01) containing the specified amendment in 96-well plates. Plates were sealed and incubated at 37°C in a benchtop microplate shaker (1,200 rpm). 2 μl of each culture was transferred into 198 μl fresh medium every 24 h. At the specified timepoints, aliquots were diluted in M9 medium and CFP and YFP values were measured. Standard curves for conversion of fluorescence to OD_600_ equivalents were generated from saturated cultures grown in tubes (for t = 0 readings only) or mono-culture controls grown in 96-well plates (from t = 1 day on). Competition experiments with pACYCDuet-1-pLac plasmids expressing *btuR* and/or *yciK* were performed similarly except that strains were pre-cultured in M9 glucose (0.2%) medium with Met.

### Whole genome sequencing and analysis

Evolved populations were grown in M9 medium supplemented with 0.35 nM pCbl, while evolved isolates and the *ΔmetE* ancestor were cultured in M9 medium supplemented with Met. Genomic DNA was isolated with a DNeasy Blood and Tissue Kit (Qiagen) and submitted to Novogene (Sacramento, CA, USA) for library preparation and whole genome sequencing using an Illumina NovaSeq 6000.

Identification of mutations was performed by Novogene by comparison to the *E. coli* MG1655 reference genome (accession PRJNA57779). SNPs and InDels were detected using SAMtools with the parameter ‘mpileup -m 2 -F 0.002 -d 1000’ and annotated using ANNOVAR (66, 67). The results were filtered such that the number of support reads for each SNP/InDel was greater than 4 and the mapping quality of each SNP/InDel was higher than 20. SVs were detected by BreakDancer and annotated by ANNOVAR (68). SVs were filtered by removing those with fewer than 2 supporting PE reads. A comparison to the *ΔmetE* ancestor was made to eliminate mutations in the genome present prior to the laboratory evolution.

### Measurement of btuB and btuR expression in evolved isolates by RT-qPCR

The *E. coli* Δ*metE* ancestor and evolved isolates S2, S3, S4, R1, and R3 were streaked onto LB agar and grown at 37°C. Three individual colonies of each strain were inoculated into M9 glycerol medium supplemented with 1 nM pCbl. After 22 hours of growth at 37°C, the cultures were passaged into fresh medium at an OD_600_ of 0.01 and grown for 20 hours. Cultures were then diluted to an OD_600_ of 0.03 in fresh medium and samples were collected during exponential phase (OD_600_ ~0.4). 875 μl of each culture was mixed with 1.75 ml RNAprotect® Bacteria Reagent and RNA was extracted with the RNeasy Mini Kit (Qiagen). On-column DNase digestion was performed to remove DNA. RNA was quantified using a Qubit fluorometer (Invitrogen). 100 ng of RNA was used to synthesize cDNA in 20 μl reactions with iScript RT Supermix (Biorad). 5 μl of 50-fold diluted cDNA was used in 20 μl reactions with SsoAdvanced Universal SYBR Green Supermix (Biorad), with reactions performed in triplicate with each primer pair (5 μM). qPCR was performed on a CFX96 Touch Real-Time PCR Detection System (Biorad) with the following conditions: one cycle of denaturation at 95°C for 3 min, and 40 cycles of denaturation at 95°C for 10 sec and extension at 59°C for 30 sec. C_q_ values were normalized against previously reported reference genes *mdh* and *rpoA* (69) and relative gene expression was calculated using the ΔΔC_q_ method. Primers used for qPCR are listed in Table S2.

### Corrinoid bioassay to assess cobamide uptake and retention

*E. coli ΔbtuR ΔmetE* and *ΔmetE* strains were pre-cultured in M9 glycerol medium supplemented with Met. The strains were then inoculated at an OD_600_ of 0.01 in 1 ml M9 glycerol medium supplemented with AdopCbl, CNpCbl, AdoCbl, or CNCbl. The medium was also supplemented with 0.02 mg/ml Met, a concentration that ensured saturating growth of the pCbl cultures but did not affect growth of the *ΔmetE* strain in the subsequent bioassay. Cultures were incubated at 37°C with aeration for 22 hours. Cultures containing adenosylcobamides were prepared under red light and incubated in the dark. 750 μl of each culture was centrifuged for 5 min at 6,000 x g to pellet cells. 600 μl of the supernatant was passed through a 0.22 μm filter. The cell pellet was washed twice with 0.85% saline and resuspended in 750 μl saline. All samples were then incubated at 100°C for 20 min. Samples containing the pellet fraction were centrifuged for 5 min at 6,000 x g and 600 μl of supernatant was removed to use as the cell lysate. The *E. coli* bioassay was performed in 96 well plates as described (65), except M9 glycerol was used as the growth medium and plates were incubated at 37°C in a microplate shaker for 22 hours prior to measurement of OD_600_. The concentration of cobamides in each sample was determined using standard curves generated with CNpCbl and CNCbl.

### Corrinoid extraction and analysis

*E. coli ΔbtuR ΔmetE* was pre-cultured in M9 glycerol medium supplemented with Met. OD_600_ was measured and cells were inoculated into 250 ml M9 glycerol medium containing 1 nM pCbl or Cbl at an OD_600_ of 0.01. The medium with 1 nM pCbl was supplemented with Met to enable growth of *E. coli ΔbtuR ΔmetE*. Cultures were grown at 37°C with aeration for 22 h. Cells were collected by centrifugation and washed twice with saline. Corrinoids were extracted with KCN as described (65). Extractions were analyzed on an Agilent Zorbax SB-Aq column (5 μm, 4.6 x 150 mm) with an Agilent 1200 series HPLC equipped with a diode array detector using Method 2 (10). Cobamides in each sample were quantified using standard curves generated with CNpCbl and CNCbl.

## Acknowledgements

We thank Alexa Nicolas for advice on whole genome sequencing and analysis, Sophia Adler for advice with RT-qPCR, Zoila Alvarez-Aponte, Markos Koutmos and members of the Taga Lab for helpful discussions, and Janani Hariharan, Dennis Suazo, and Eleanor Wang for critical reading of the manuscript. We are grateful to Olga Sokolovskaya for synthesis of AdopCbl and Sebastian Gude for construction of pSG013 and pSG015. This work was funded by National Institutes of Health grants R35GM139633 to M.E.T. and 5K99GM143653-02 to Z.F.H. R.R.P. was supported by the NIH T32GM132022 training grant.

## Notes

### Competing Interest Statement

The authors have declared no competing interest.

### Summary of Updates

This version of the manuscript has been revised with updates to the text (including additional data) and supplemental files.

