## Supplemental Figures for "Laboratory evolution of *E. coli* with a natural vitamin B_12_ analog reveals roles for cobamide uptake and adenosylation in methionine synthase-dependent growth"

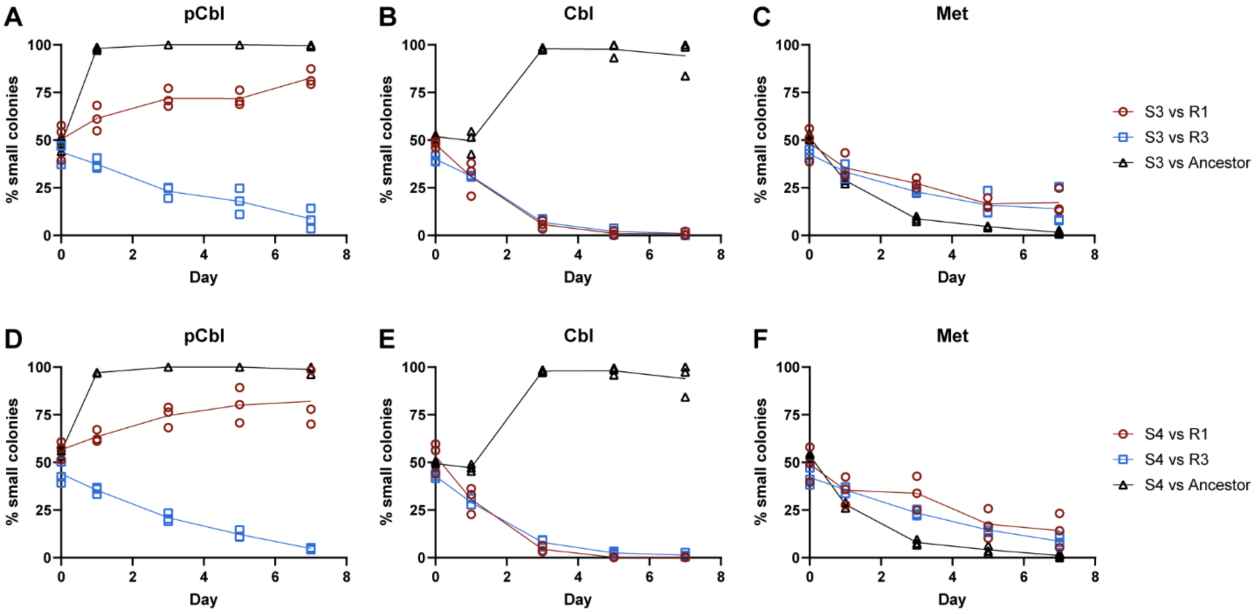


**Figure S1. Growth competition of small and regular sized isolates from Culture 8.** Isolates S3 (A-C) and S4 (D-F) were competed in co-culture against isolates R1 and R3 and the ancestor strain with daily passaging in medium containing either 0.35 nM pCbl, 0.35 nM Cbl, or 0.1 mg/ml Met. Cultures were diluted and plated on the indicated days to quantify the fraction of small colonies (S3 or S4). Lines connect the means of three biological replicates.

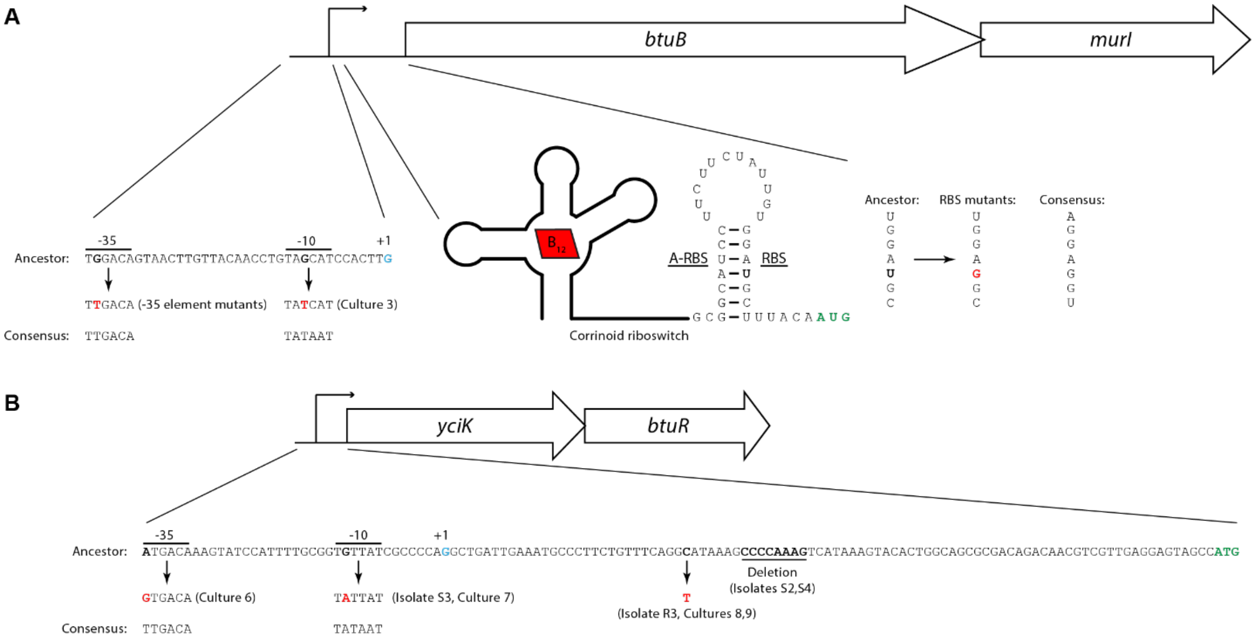


**Figure S2. Mutations in the *btuB-murI* and *yciK-btuR* operons in evolved isolates and populations.** Nucleotides in the native sequence that were mutated are bolded, with their corresponding changes shown in red. Transcriptional start sites and start codons are shown in blue and green, respectively. The consensus sequences for the $\sigma^{70}$ promoter -35 element, -10 element, and RBS are shown for comparison. The promoter for the *yciK-btuR* operon has not been experimentally characterized and was predicted by PromoterHunter. A) Changes in the promoter and RBS/$B_{12}$ riboswitch of the *btuB-murI* operon likely cause increased expression. The mutations found in the -35 element, -10 element, and RBS result in sequences closer to their respective consensus sequences. Additionally, the mutation in the RBS may disrupt its translation-inhibiting interaction with the anti-RBS (A-RBS) within the corrinoid riboswitch. B) Changes to the promoter and 5′ UTR of the *yciK-btuR* operon. *yciK-btuR* is on the minus strand of the chromosome and the reverse complement of the promoter and 5′ UTR is shown. The mutation found in the -10 element likely increases transcription, as the sequence is closer to the consensus sequence. It is unclear how the other mutations affect expression of the operon.

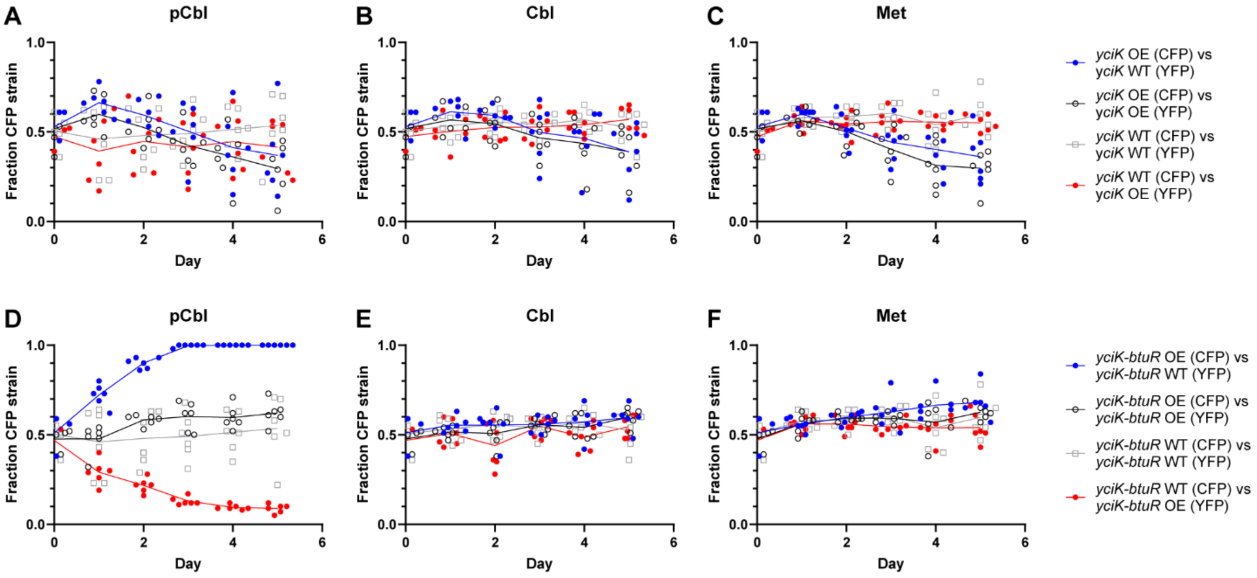


**Figure S3. Overexpression of *yciK* does not confer a growth advantage with pCbl.** CFP- and YFP-expressing Δ*metE* strains overexpressing *yciK* (OE) or producing native levels of *yciK* (WT) (A-C), and Δ*metE* strains overexpressing *yciK-btuR* (OE) or producing native levels of *yciK-btuR* (WT) (D-F), were competed in co-culture with daily passaging for five days in medium containing either 1 nM pCbl, 1 nM Cbl, or 0.1 mg/ml Met. The fraction of the CFP-expressing strain in each co-culture is plotted. Control co-cultures containing CFP- and YFP-expressing strains in the same genetic background (black and gray) were included to rule out a growth disadvantage caused by either fluorescent protein. Lines connect the means of six biological replicates.

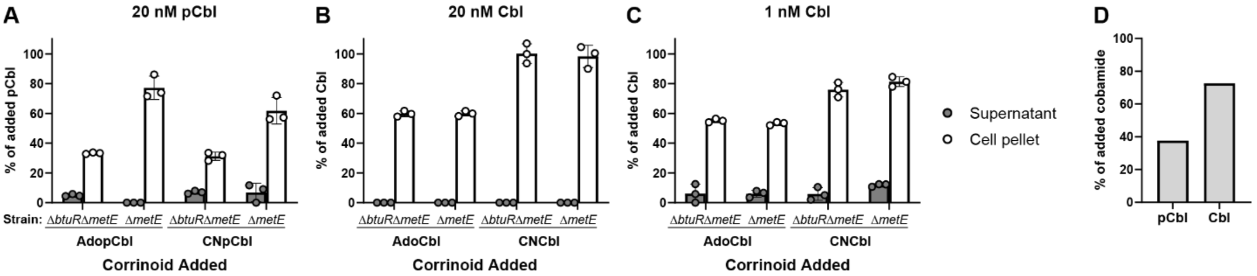


**Figure S4. Uptake and retention of cobamides by Δ*btuR* Δ*metE* and Δ*metE* strains.** A-C) The total corrinoid concentration in supernatants and lysed cells of *E. coli* Δ*btuR* Δ*metE* and Δ*metE* strains grown with 20 nM AdopCbl or CNpCbl (A), 20 nM AdoCbl or CNCbl (B), and 1 nM AdoCbl and CNCbl (C) was measured with an *E. coli* bioassay (65). The cultures were supplemented with 0.02 mg/ml Met to ensure that those containing pCbl grew to saturation. The calculated corrinoid concentrations were normalized to the amount of cobamide added to the culture. Data represent the average and standard deviation of three biological replicates. D) Corrinoid extractions of *E. coli* Δ*btuR* Δ*metE* cells grown with 1 nM CNpCbl or CNCbl were analyzed by HPLC to quantify the amount of cobamide present in the cell pellets. The cultures were supplemented with 0.1 mg/ml Met to ensure the one containing pCbl grew to saturation.
