## Supplemental Table 2 for "Laboratory evolution of *E. coli* with a natural vitamin B_12_ analog reveals roles for cobamide uptake and adenosylation in methionine synthase-dependent growth"

**Table S2. qPCR primers**

| **Primer** | **Sequence** | **Primer efficiency** |
| --- | --- | --- |
| mdh-F | CGGTTATTGGCGGTCACTCT | 95.8% (r^2^ = 0.997) |
| mdh-R | CGTTCTGGATGCGTTTGGTC |  |
| rpoA-F | GGAAGAAGATGAGCGCCCAA | 99.2% (r^2^ = 0.999) |
| rpoA-R | CGCGCTGCTTCAACATTGTA |  |
| btuB-F | TGGTCGTTATGATTCGTCGG | 98.0% (r^2^ = 0.999) |
| btuB-R | GTCGTAGTCTGTTTCTGCCA |  |
| btuR-F | TCAGCAGCGAGTGAAAGAAA | 99.8% (r^2^ = 0.993) |
| btuR-R | TTTTGCCTTTTCCATTGCCG |  |
